# Resolving phylogeny and polyploid parentage using genus-wide genome-wide sequence data from birch trees

**DOI:** 10.1101/2020.07.13.200444

**Authors:** Nian Wang, Laura J. Kelly, Hugh A. McAllister, Jasmin Zohren, Richard J. A. Buggs

## Abstract

Numerous plant genera have a history including frequent hybridisation and polyploidisation, which often means that their phylogenies are not yet fully resolved. The genus *Betula*, which contains many ecologically important allopolyploid tree species, is a case in point. We generated genome-wide sequence data for 27 diploid and 31 polyploid *Betula* species or subspecies using restriction site associated DNA (RAD) sequences assembled into contigs with a mean length of 675 bp. We reconstructed the evolutionary relationships among diploid *Betula* species using both supermatrix and species tree methods. We identified progenitors of the polyploids according to the relative rates at which their reads mapped to contigs from different diploid species. We sorted the polyploid reads into different putative sub-genomes and used the extracted contigs, along with the diploid sequences, to build new phylogenies that included the polyploid sub-genomes. This approach yielded a highly evidenced phylogenetic hypothesis for the genus *Betula*, including the complex reticulate origins of the majority of its polyploid taxa. The genus was split into two well supported clades, which differ in their seed-wing morphology. We propose a new taxonomy for *Betula*, splitting it into two subgenera. We have resolved the parentage of many widespread and economically important polyploid tree species, opening the way for their population genomic study.

## 1. Introduction

The evolution of plant diversity cannot be fully understood unless we can reconstruct evolutionary relationships for allopolyploids (hybrid species with duplicated genomes). While phylogenomic approaches that use thousands of loci can resolve the phylogenies of diploid taxa with a history of hybridization (Folk et al., 2017; Fontaine et al., 2015; Li et al., 2016), allopolyploids, which are common in plants (Barker et al., 2016), remain hard to place in phylogenetic trees (Oxelman et al., 2017). When molecular markers are sequenced from polyploids, it is difficult to phase them into their parental subgenomes (Eriksson et al., 2018), and it is easy to mistake homoeologs (genes duplicated in allopolyploidisation events) for paralogs (genes arising from duplication within a genome) and vice versa (Brysting et al., 2011; Linder and Rieseberg, 2004). Approaches to resolving the phylogenetic origins of tetraploids (but not higher ploidy levels) have determined parental genomes using heuristic methods (Jones et al., 2013; Lott et al., 2009) or long-read sequences (Rothfels et al., 2017). While phylogenomic approaches are sometimes used to detect the presence of hybrid polyploids (McKain et al., 2016; Morales-Briones et al., 2018), we are not aware of studies that have used phylogenomic data to resolve polyploid origins.

Resolving parental origins of polyploid subgenomes unlocks progress in their genomic characterisation. Knowledge of the origins of the allohexaploid genome of *Triticum aestivum* (bread wheat) allowed further characterisation to be assisted by sequencing of the close relatives of its parents: *Aegilops tauschii* and *Triticum turgidum* ssp. *dicoccoides* (Avni et al., 2017; Luo et al., 2017). Similarly, the allotetraploid genome of *Gossypium hirsutum* (Upland cotton) has been illuminated by sequencing of two close diploid relatives of its progenitor genomes (Li et al., 2015). However, for many polyploids of economic and ecological importance, we do not know the identity of the closest living relatives of their progenitor genomes. We need accessible phylogenomic approaches to make this possible.

A relatively inexpensive method of generating genome-wide marker data for phylogenomics from large numbers of individuals is through sequencing of restriction-site associated DNA (RAD) libraries with short reads of 50-150 bp. This is widely used as a method of genome-wide SNP genotyping in non-model organisms for population genomic analyses (Andrews et al., 2016; Barchi et al., 2011; Emerson et al., 2010; Etter et al., 2011; Hohenlohe et al., 2010; Zohren et al., 2016). SNP data from RAD-seq has also been used in phylogenetic reconstruction using supermatrix approaches which assume that all loci have the same evolutionary history (Cariou et al., 2013; Cruaud et al., 2014; Eaton and Ree, 2013; Eaton et al., 2016; Gonen et al., 2015; Hipp et al., 2014; Massatti et al., 2016; Pante et al., 2015; Rubin et al., 2012; Wagner et al., 2013). A few studies have used species tree approaches, which take into account the possibility of different evolutionary histories for separate loci, for analysis of short read RAD-seq data. For example, Eaton and Ree (2013) used RAD loci inferred from single end reads to build a species tree in the genus *Pedicularis* (lousewort) (Eaton and Ree, 2013). DaCosta and Sorensen (2016) used single end reads to construct species trees in two avian genera and Hou et al. (2015) used paired-end reads to build a species tree for the genus *Diapensia* (pincushion plant) (DaCosta and Sorenson, 2016; Hou et al., 2015).

We reasoned that extra power for phylogenetic analysis may be gained by sequencing RAD libraries with 300 bp paired-end reads and assembling these reads against a reference genome to generate longer contigs spanning restriction enzyme and variable sites. These contigs can be aligned to each other and individual phylogenies reconstructed for each locus, for input into species tree methods, or the alignments combined, for a supermatrix approach. So far, we are not aware of any studies which have sequenced longer RAD loci in an attempt to gain greater power for species tree methods.

The genus *Betula* (birches) includes about 65 species and subspecies with ranges across the Northern Hemisphere (Ashburner and McAllister, 2016). Some act as keystone species of forests across Eurasia and North America (Ashburner and McAllister, 2016). Various birch species are planted for timber, paper, carbon sequestration and ecological restoration, but some birch species are endangered with narrow distributions and there is concern about the increasing threat posed by the bronze birch borer (Muilenburg and Herms, 2012; Shaw et al., 2014). Previous phylogenetic analyses of *Betula* using nuclear genes (ITS, NIA and ADH), chloroplast genes (matK and rbcL) and AFLPs provided limited resolution of relationships among species and partly contradicted each other (Järvinen et al., 2004; Li et al., 2005; Li et al., 2007; Schenk et al., 2008). In addition, molecular phylogenies based on nuclear genes contradict some species groupings proposed in a recent monograph based on morphology, such as the placement of the ecologically and economically important *B. maximowicziana* (monarch birch) (Ashburner and McAllister, 2016; Wang et al., 2016).

Hybridisation is frequent and has been extensively documented in *Betula* (Anamthawat-Jónsson and Tómasson, 1990; Anamthawat-Jónsson and Tómasson, 1999; Anamthawat-Jónsson and Thórsson, 2003; Anamthawat-Jónsson et al., 2010; Barnes et al., 1974; Eidesen et al., 2015; Johnsson, 1945; Tsuda et al., 2017; Wang et al., 2014). Polyploidy is also common within *Betula*, with nearly 60% of species being polyploids (Wang et al., 2016) and ploidy ranging from diploid to dodecaploid (Ashburner and McAllister, 2016). Some species contain different cytotypes, such as *B. chinensis* (6x and 8x) (Ashburner and McAllister, 2016). The origins of most polyploids in the genus are unresolved. One of the best studied is the tetraploid *B. pubescens* (downy birch), with different lines of evidence suggesting as candidate parents: *B. pendula* based on RAPD markers (Howland et al., 1995), *B. humilis* or *B. nana* based on ADH (Järvinen et al., 2004), *B. humilis* based on morphology (Walters, 1968) or *B. lenta* based on SNPs (Salojarvi et al., 2017). This uncertainty hinders genomic research on *B. pubescens*, the most widespread birch tree in Europe and western Asia.

Here, in order to better resolve the phylogeny of *Betula* and elucidate the parental origins of its polyploid species we use a RAD-seq approach with reads assembled against the *B. pendula* reference genome (Salojarvi et al., 2017). We construct the phylogeny of diploid species using supermatrix and species tree methods. As a heuristic method for analyzing the origins of the polyploid species, we create a reference using contigs from all diploid species and compare the genomic similarity between each polyploid species and all diploids by mapping reads of each polyploid species to the reference. Polyploid taxa should have a higher level of genetic similarity to diploids closely related to their ancestors and hence a higher number of mapped reads. These approaches together yielded a well-resolved history for the genus *Betula*, including polyploid taxa.

## 2. Materials and Methods

### 2.1. Sample collection

Samples were obtained from living collections in Stone Lane Gardens (SL hereafter), Ness Gardens (N hereafter), the Royal Botanic Garden Edinburgh (RBGE) or collected from the wild by the research group (Table S1). The genome size of most of these taxa has been obtained (Wang et al., 2016) and morphological characters were used to confirm the identity of each taxon sampled. *Alnus inokumae* was chosen as the outgroup as *Alnus* has been shown to be sister to *Betula* (Li et al., 2007). In addition, *A. orientalis* and *Corylus avellana* were included for marker development. Herbarium specimens of most of these samples have been deposited at the Natural History Museum London and RBGE with accession numbers provided in Table S1.

### 2.2. DNA extraction, RAD library preparation and Illumina sequencing

Genomic DNA was isolated from silica-dried cambial tissue or leaves following a modified 2 X CTAB (cetyltrimethylammonium bromide) protocol (Wang et al., 2013). The isolated DNA was assessed with a 1.0% agarose gel and measured with a Qubit 2.0 Fluorometer (Invitrogen, Life technologies) using Broad-range assay reagents. RAD libraries were prepared following the protocol of Etter et al. (2011) with slight modifications (Etter et al., 2011). Briefly, 0.5-1.0 μg of genomic DNA for each sample was heated at 65°C for 2-3 hours prior to digestion with PstI (New England Biolabs, UK). This enzyme has a 6 bp recognition site and leaves a 4 bp overhang. Digestion was followed by ligation of barcoded P1 adapters. Ligated DNA was sheared using a Bioruptor (KBiosciences, UK) instrument in 1.5 mL tubes (high intensity, duration 30 s followed by a 30 s pause which was repeated eight times). Sheared fragments were evenly distributed between 100 bp and 1500 bp and fragments of ~600 bp were selected using Agencourt AMPure XP Beads (New England Biolabs) following a protocol of double-size selection. Briefly, a ratio of bead:DNA solution of 0.55 was used to remove large fragments and then a second round of size selection was conducted, using 5 μl of bead solution concentrated from a starting volume of 20 μl. After end-repair and A-tailing, the size-selected DNA was ligated to P2 adapters (400 nm) and PCR amplified. PCR amplification was carried out in 25 μl reactions consisting of 0.46 vol ddH_2_O and template DNA (4-5 ng), 0.5 vol 2×Phusion Master Mix (New England Biolabs), and 0.04 vol P1 and P2 amplification primers (10 nm), using the following cycling parameters: 98°C for 30 s followed by 12 cycles of 98°C for 10 s and 72°C for 60 s. Three or four independent PCR replicates were conducted for each sample to achieve a sufficient amount of the library. The final library was quantified using a Bioanalyzer and a Qubit 2.0 Fluorometer (Invitrogen, Life Technologies) using high-sensitivity assay reagents and was normalized prior to sequencing. The quantified library was sequenced on an Illumina MiSeq machine using MiSeq Reagent Kit v3 (Illumina) at the Genome Centre of Queen Mary University of London.

### 2.3. RAD data trimming and demultiplexing

The raw data were trimmed using Trimmomatic (Bolger et al., 2014) in paired-end mode with the following steps. First, LEADING and TRAILING steps were used to remove bases from the ends of a read if the quality is below 20. Then a SLIDINGWINDOW step was performed with a window size of 1 and a required quality of 20. Finally, a MINLENGTH step was used to discard reads shorter than 100bp. FastQC was used to check various parameters of sequence quality in both raw and trimmed datasets (Andrews, 2014). The trimmed data were demultiplexed, using the process_radtags module of Stacks (Catchen et al., 2013).

### 2.4. Reads mapping, sequence alignment and trimming

The whole genome assembly of *B. pendula* (Salojarvi et al., 2017) was used as a reference for mapping our RAD data, to separate orthologous loci (i.e. mapped segments of DNA) from paralogous loci (Wang et al., 2013), and to anchor reads with adjacent restriction cutting sites. Mapping of trimmed reads for each sample was conducted using the ‘Map Reads to Reference’ tool in the CLC Genomics Workbench v. 8. A similarity value of 0.8 and the fraction value of 0.8 were applied as the threshold. Reads with non-specific matches were discarded and any regions with coverage of below three were removed. A consensus sequence with a minimum contig length of 300bp was created for each sample. *Betula glandulosa* was excluded from further analysis because only 216 loci were mapped at a sufficient read depth. Multiple sequence alignments for individual loci were generated using mafft v.6.903 (Katoh et al., 2005) with default parameters. Aligned sequences were trimmed using trimAl v1.2rev59 (Capella-Gutierrez et al., 2009); gaps present in 40% of taxa or above were removed (-gt 0.6).

### 2.5. Species tree inference

Two datasets were used for phylogenetic analysis: dataset 1 (D1 hereafter) includes 20 diploid *Betula* samples and dataset 2 (D2 hereafter) 27 diploid samples. In D2, some species were represented by more than one sample (Table S2). RAD loci ≥ 300bp in length that occurred in a minimum of four *Betula* samples were used for gene tree inference. The gene tree for each locus was estimated using the maximum-likelihood method (ML) in RAxML v. 8.1.16 (Stamatakis, 2006). A rapid bootstrap analysis with 100 bootstraps and 10 searches was performed under a GTR+GAMMA nucleotide substitution model. The species tree was estimated from the gene trees with ASTRAL-II v5.5.7 (Mirarab and Warnow, 2015) and ASTRID (Vachaspati and Warnow, 2015). Branch support in the ASTRAL and ASTRID trees was assessed via calculation of local posterior probabilities based on the gene tree quartet frequencies (Sayyari and Mirarab, 2016) and bootstrapped gene trees (Vachaspati and Warnow, 2015), respectively. All loci used for building gene trees and inferring the species trees were concatenated into a supermatrix, using custom shell scripts, which was analysed in RAxML v. 8.1.16 using the same settings as above. The consensus tree generated above was visualised in FigTree v.1.3.1 (http://tree.bio.ed.ac.uk/software/figtree).

### 2.6. Phylogenetic Networks

We used the Species Networks applying Quartets (SNaQ) method (Solís-Lemus and Ané, 2016) implemented in the software PhyloNetworks 0.5.0 (Solis-Lemus et al., 2017) to investigate whether the species tree without hybridisation events or a phylogenetic network with one or more hybridisation events better describes the diploid species relationships within *Betula*. Phylogenetic trees generated in RAxML were used to estimate quartet concordance factors (CFs), which represent the proportion of genes supporting each possible relationship between each set of four species. These CFs were then used to reconstruct phylogenetic networks under incomplete lineage sorting (ILS) and differing numbers of hybridisation events, and to calculate their respective pseudolikelihoods. To determine whether a tree accounting for ILS or a network better fits the observed data, we estimated the best phylogenetic network with hybridisation events (h) ranging from 0 to 5 using the phylogeny obtained with ASTRID as a starting tree. Using a value of h=0 will yield a tree without reticulation, h=1 will yield a network with a maximum of one reticulation and so forth. The fit of trees and networks to the data was evaluated based on pseudo-deviance values, and estimated inheritance probabilities (i.e. the proportion of genes contributed by each parental population to a hybrid taxon) were visualised. This test compares the score of each network based on the negative log-pL, where the network with the lowest value has the best fit.

### 2.7. Identification of putative diploid progenitors of polyploid species

We sought to infer the putative origins of polyploid species of *Betula* by read-mapping. First, we used RAD loci present in at least 15 out of 20 diploid *Betula* taxa to create a reference; the reference comprised 88,488 sequences in total, representing 5,045 loci. All the sequences were concatenated and separated with lengths of 500 Ns. Trimmed reads of polyploid taxa were mapped individually to this single reference containing loci from all diploid taxa using strict parameters in CLC Genomic Workbench: a fraction value of 0.9 and a similarity value of at least 0.995. Reads with non-specific matches were discarded. Any region with a coverage of below three was removed and the consensus sequences for each polyploid with a minimum length of 300 bp were extracted. Variable sites were represented by ambiguity codes in the consensus. Given the fact that the number of loci available for each diploid species was variable (Table S2), we plotted the proportion of loci in the reference for each diploid species to which reads from each polyploid species mapped. We expected a higher proportion of consensus loci to be mapped in those diploids that were progenitor species of each polyploid species. We therefore sought to identify diploid progenitors for each polyploid based on the number of mapped consensus loci assuming that the number of progenitors could not be more than half the ploidy level of each polyploid. In addition, we mapped reads from diploid species to the reference containing loci from all diploid taxa using the same parameters as described above. We found a small number of loci of each diploid were mapped to by reads from other diploid species, with the exception of *B. calcicola* and *B. potaninii*, for which a relatively high number of loci (>1000) were mapped in each species by reads from the other (Fig. S1).

### 2.8. Phylogeny incorporating polyploid species

For those polyploids for which we could identify putative diploid parental species, we separated their homoeologues for each RAD-locus using another concatenated single reference, similar to the one described in the paragraph above, but this time containing all 50,870 loci present in a minimum of four *Betula* taxa of D1. For each of these polyploids, we extracted a RAD consensus sequence from each mapped diploid locus, with a minimum length of 200 bp; we excluded any sequence where the polyploid had not mapped to their putative parents for that locus. We then excluded loci where reads from one diploid had mapped to another diploid species (see above - Identification of putative diploid progenitors of polyploid species). We constructed phylogenies including the phased polyploid RAD loci with the diploid RAD loci (Fig. S2). Individual gene trees were constructed in RAxML v. 8.1.16 using the same parameters as described above (see - Species tree inference) and the species tree was inferred using ASTRID. The putative diploid progenitors which we included for phylogenetic analysis are provided in Table S2.

### 2.9. Simple sequence repeat analysis

To develop markers for future use in all *Betula* species for population genetic analyses, mapped consensus sequences with length equal to or greater than 300 bp were mined for simple sequence repeats (SSR) using the QDD pipeline version 3.1.2 (Meglécz et al., 2014). Consensus sequences with a repeat motif of 2–5 bp, and repeated a minimum of five times, were screened using the Downstream QDD pipeline version 3.1.2. Primer pairs were designed within 200 bp flanking regions using PRIMER3 software (Untergasser et al., 2012). The primer table output by the QDD version 3.1.2 pipeline allows selection of the best primer pair design for each SSR locus. We filtered primer pairs according to parameters provided by QDD version 3.1.2. The selected SSR loci had: a minimum number of 7 motif repeats within the SSR sequence; a maximum primer alignment score of 5; a minimum of 20 bp forward and reverse flanking region between SSR and primer sequences; and a high-quality primer design without homopolymer, nanosatellite and microsatellite sequence in the primer or flanking sequences. For polyploid species of *Betula*, *A. inokumae*, *A. orientalis* and *C. avellana*, we selected SSR loci with a minimum number of 5 motif repeats as a majority of loci had 5 or 6 motif repeats within the SSR sequence.

## 3. Results

### 3.1. RAD data description

The number of trimmed reads per diploid taxon ranges from 1,065,196 to 2,560,486 (average of 1,508,904) with between 881,333 and 2,252,171 (80.60% - 90.75%) mapped to the *B. pendula* genome for each of the 27 diploid *Betula* and 707,914 (51.64%) for the outgroup *A. inokumae* (Table S1). In D1, 162 loci are present in all 20 *Betula* diploid taxa and 7,002 present in only four of these (Fig. 1A), whereas for D2 99 loci are present in all 27 *Betula* diploid individuals and 6,078 present in only four (Fig. 1B). Contigs of ≥ 300bp, with an average length of 580.8bp - 755.8bp, varied in number between 13,597 in *A. inokumae* and 30,717 in *B. pendula* (Fig. 1C, D).

**Figure 1.**
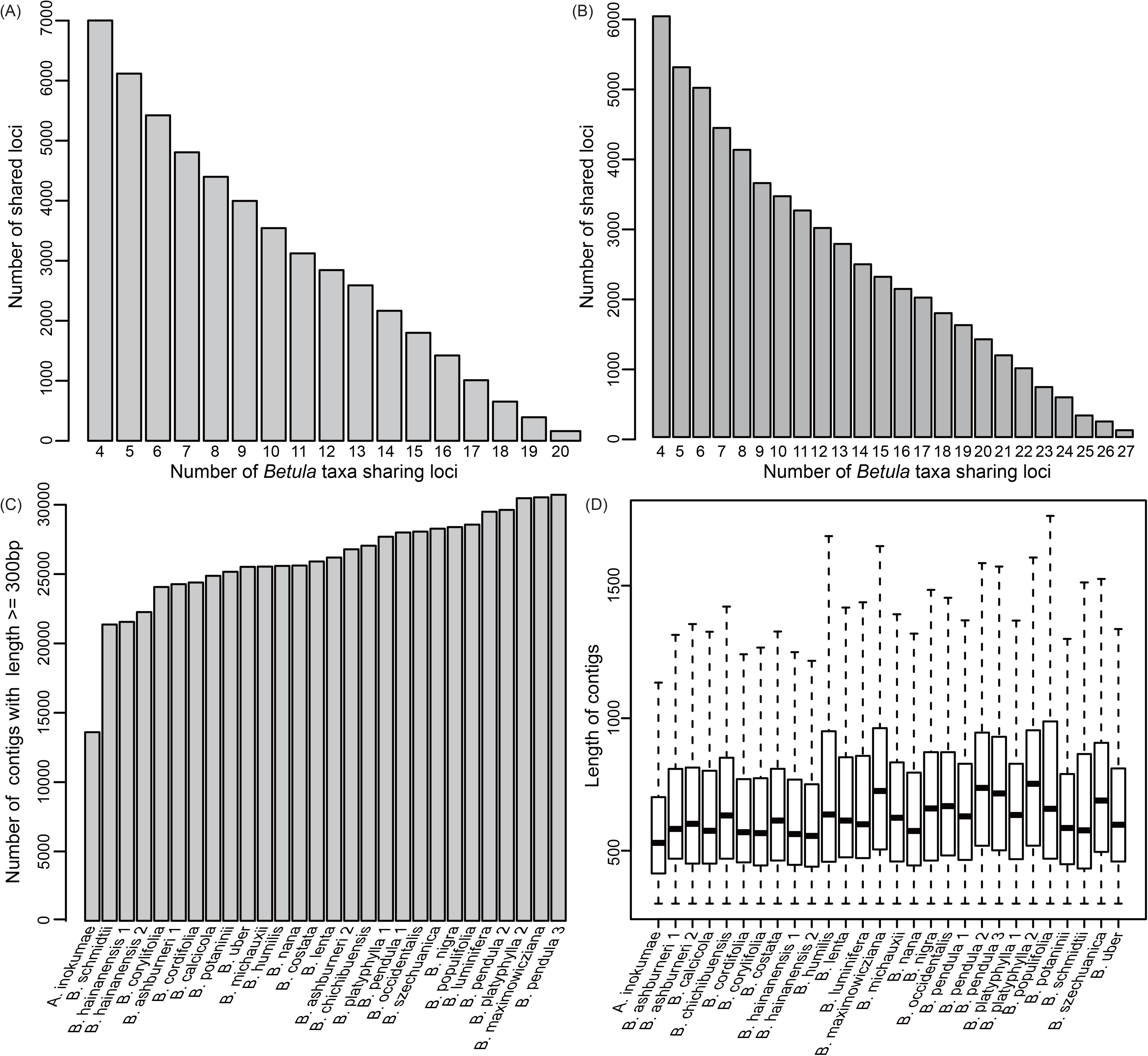
Detailed information of number of shared loci and number of contigs (A) number of shared loci only in between four and 20 of the diploid *Betula* species of D1; (B) number of shared loci only in between four and 27 of the diploid *Betula* species of D2; (C) number of contigs with length above 300bp for *A. inokumae* and each of the 27 diploid *Betula* species of D2; (D) length of contigs for *A. inokumae* and each of the 27 diploid *Betula* species of D2. The whiskers of the boxplot from the bottom to the top indicate the minimum, the first quartile, the median, the third quartile and the maximum value of contig length excluding outliers.

### 3.2. Phylogenetic inference

The concatenated D1 (50,870 loci) and D2 (58,442 loci) datasets include 31,815,738 and 35,859,769 nucleotides with 60.25% and 63.12% missing data (gaps and undetermined characters), respectively. The three approaches used for phylogenetic analysis of D1 (ASTRAL, ASTRID and supermatrix) all produced well resolved trees that split the genus into two major clades. The ASTRAL species tree (Fig. 2A) and concatenation tree (Fig. 2B) have identical topologies, whereas the ASTRID tree for this dataset differs in the placement of *B. cordifolia* (Fig. S3). Phylogenetic trees for D2 inferred with the species tree methods also separate the genus into two major clades, similar to the D1 trees, but with some differences in the placement of a small number of taxa within the largest clade (Fig. S4-S5). The concatenation tree of D2 does not recover the same major clades as the other analyses, although these differences are not well supported (Fig. S6).

**Figure 2.**
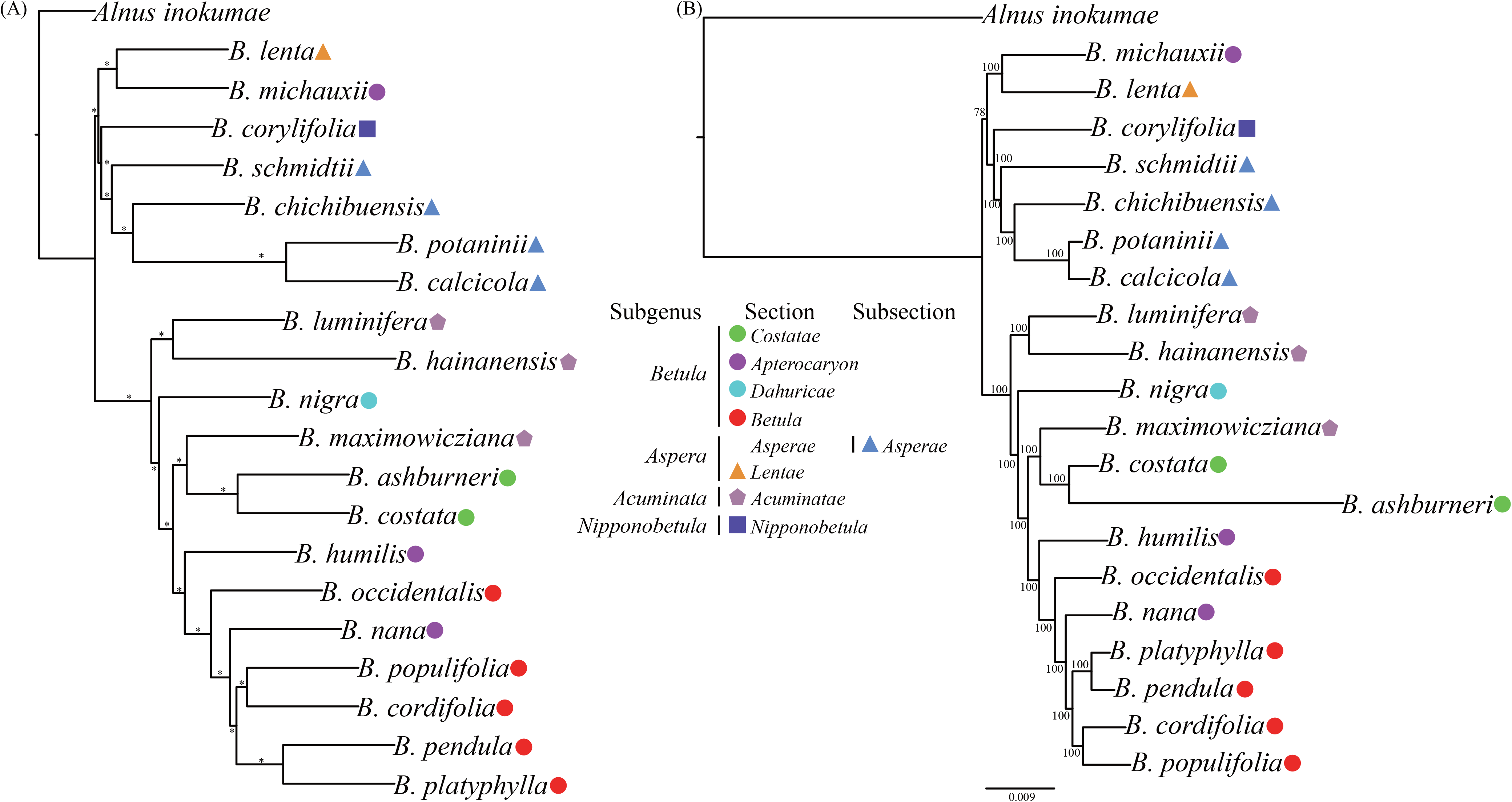
Species tree from the maximum likelihood analysis of the 20 *Betula* diploids of D1 using ASTRAL (A) and the supermatrix (B) approach based on data from 50,870 loci. Asterisks on the branches of (A) indicate local posterior probabilities of 1 and numbers on the branches of (B) are bootstrap support values. The scale bar below (B) indicates the mean number of nucleotide substitutions per site. Species were classified according to Ashburner and McAllister (2016).

### 3.3. Phylogenetic Networks

The pseudolikelihood values of hybrid nodes decreased sharply from h = 0 to h = 2, with only marginal improvements when further increasing the number of hybridisation events (Fig. S7), suggesting the best-fitting phylogenetic model is one involving two main hybridisation events. The D1 phylogenetic network when h = 2 is similar to the phylogenetic trees for this dataset (Fig. 3), but with evidence for hybridisation events involving four separate lineages within the largest of the major clades.

**Figure 3.**
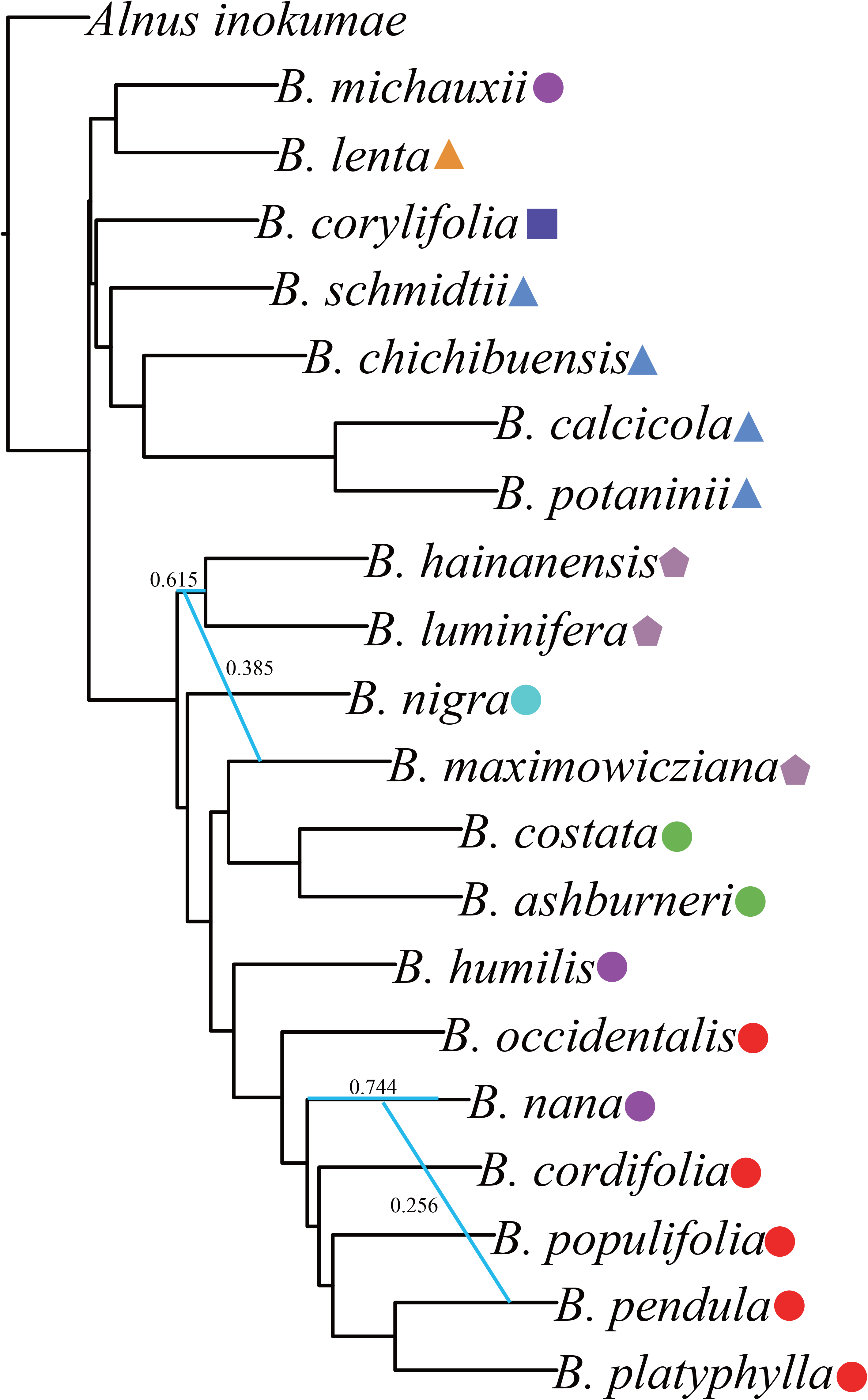
Best network inferred from SNaQ analysis of the 20 *Betula* diploids of D1 with the number of hybridization events h=2. Blue lines indicate hybrid edges and values beside the blue line indicate estimated inheritance probabilities.

### 3.4. Read-mapping of polyploid species

For 28 of the polyploid species or varieties (80%) the mapping analysis identified two or more parental lineages (Fig. 4), represented by 17 diploid species (Table 1). Five polyploids, all with the ploidy level ≥ 8x, have putative progenitors from both of the two major diploid clades. *Betula nigra* seems not to be the putative progenitor of any polyploid species whereas *B. humilis* represents the putative parental lineage of up to nine polyploids (Table 1).

**Table 1.**
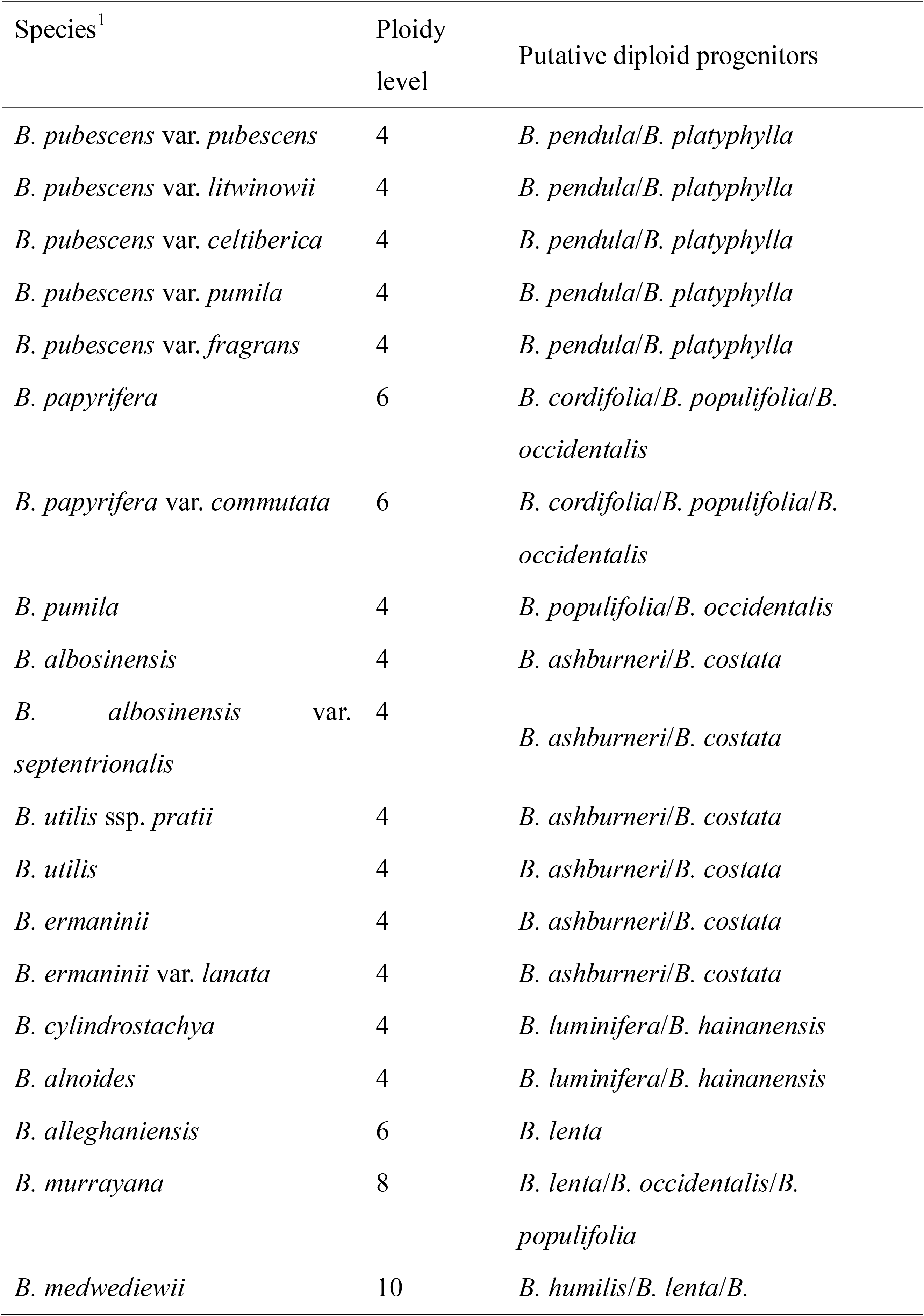
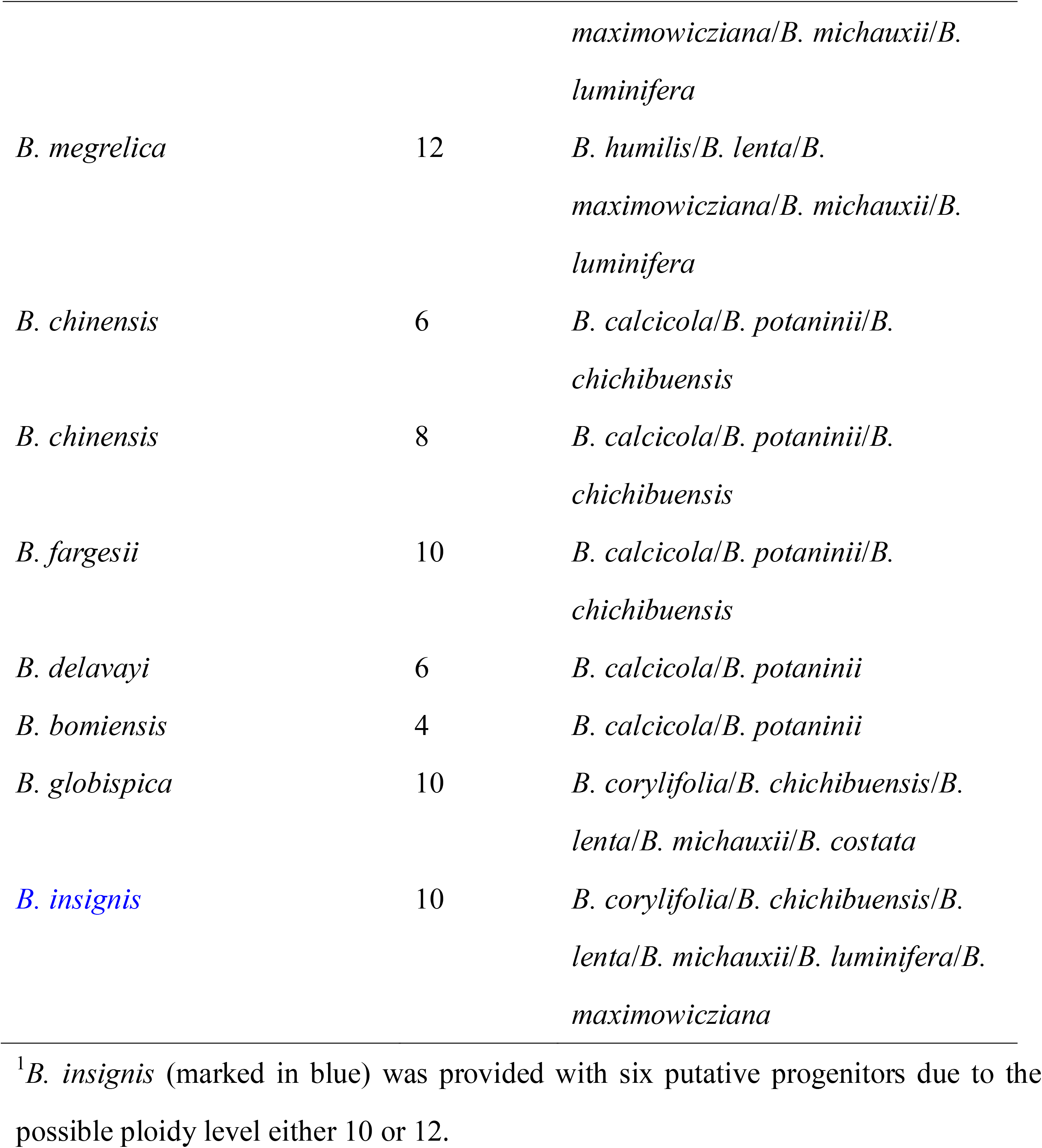
Putative diploid progenitors suggested for polyploids of *Betula* and included for phylogenetic analysis.

| Species<sup>1</sup> | Ploidy<br>level | Putative diploid progenitors |
| --- | --- | --- |
| *B. pubescens* var. *pubescens* | 4 | *B. pendula*/*B. platyphylla* |
| *B. pubescens* var. *litwinowii* | 4 | *B. pendula*/*B. platyphylla* |
| *B. pubescens* var. *celtibérica* | 4 | *B. pendula*/*B. platyphylla* |
| *B. pubescens* var. *pumila* | 4 | *B. pendula*/*B. platyphylla* |
| *B. pubescens* var. *fragrans* | 4 | *B. pendula*/*B. platyphylla* |
| *B. papyrifera* | 6 | *B. cordifolia*/*B. populifolia*/*B. occidentalis* |
| *B. papyrifera* var. *commutata* | 6 | *B. cordifolia*/*B. populifolia*/*B. occidentalis* |
| *B. pumila* | 4 | *B. populifolia*/*B. occidentalis* |
| *B. albosinensis* | 4 | *B. ashburneri*/*B. costata* |
| *B. albosinensis* var. *septentrionalis* | 4 | *B. ashburneri*/*B. costata* |
| *B. utilis* ssp. *pratii* | 4 | *B. ashburneri*/*B. costata* |
| *B. utilis* | 4 | *B. ashburneri*/*B. costata* |
| *B. ermaninii* | 4 | *B. ashburneri*/*B. costata* |
| *B. ermaninii* var. *lanata* | 4 | *B. ashburneri*/*B. costata* |
| *B. cylindrostachya* | 4 | *B. luminifera*/*B. hainanensis* |
| *B. alnoides* | 4 | *B. luminifera*/*B. hainanensis* |
| *B. alleghaniensis* | 6 | *B. lenta* |
| *B. murrayana* | 8 | *B. lenta*/*B. occidentalis*/*B. populifolia* |
| *B. medwediewii* | 10 | *B. humilis*/*B. lenta*/*B.* |
|  |  | *maximowicziana/B. michauxii/B. luminifera* |
| *B. megrelica* | 12 | *B. humilis/B. lenta/B. maximowicziana/B. michauxii/B. luminifera* |
| *B. chinensis* | 6 | *B. calcicola/B. potaninii/B. chichibuensis* |
| *B. chinensis* | 8 | *B. calcicola/B. potaninii/B. chichibuensis* |
| *B. fargesii* | 10 | *B. calcicola/B. potaninii/B. chichibuensis* |
| *B. delavayi* | 6 | *B. calcicola/B. potaninii* |
| *B. bomiensis* | 4 | *B. calcicola/B. potaninii* |
| *B. globispica* | 10 | *B. corylifolia/B. chichibuensis/B. lenta/B. michauxii/B. costata* |
| *B. insignis* | 10 | *B. corylifolia/B. chichibuensis/B. lenta/B. michauxii/B. luminifera/B. maximowicziana* |

<sup>1</sup>*B. insignis* (marked in blue) was provided with six putative progenitors due to the possible ploidy level either 10 or 12.

**Figure 4.**
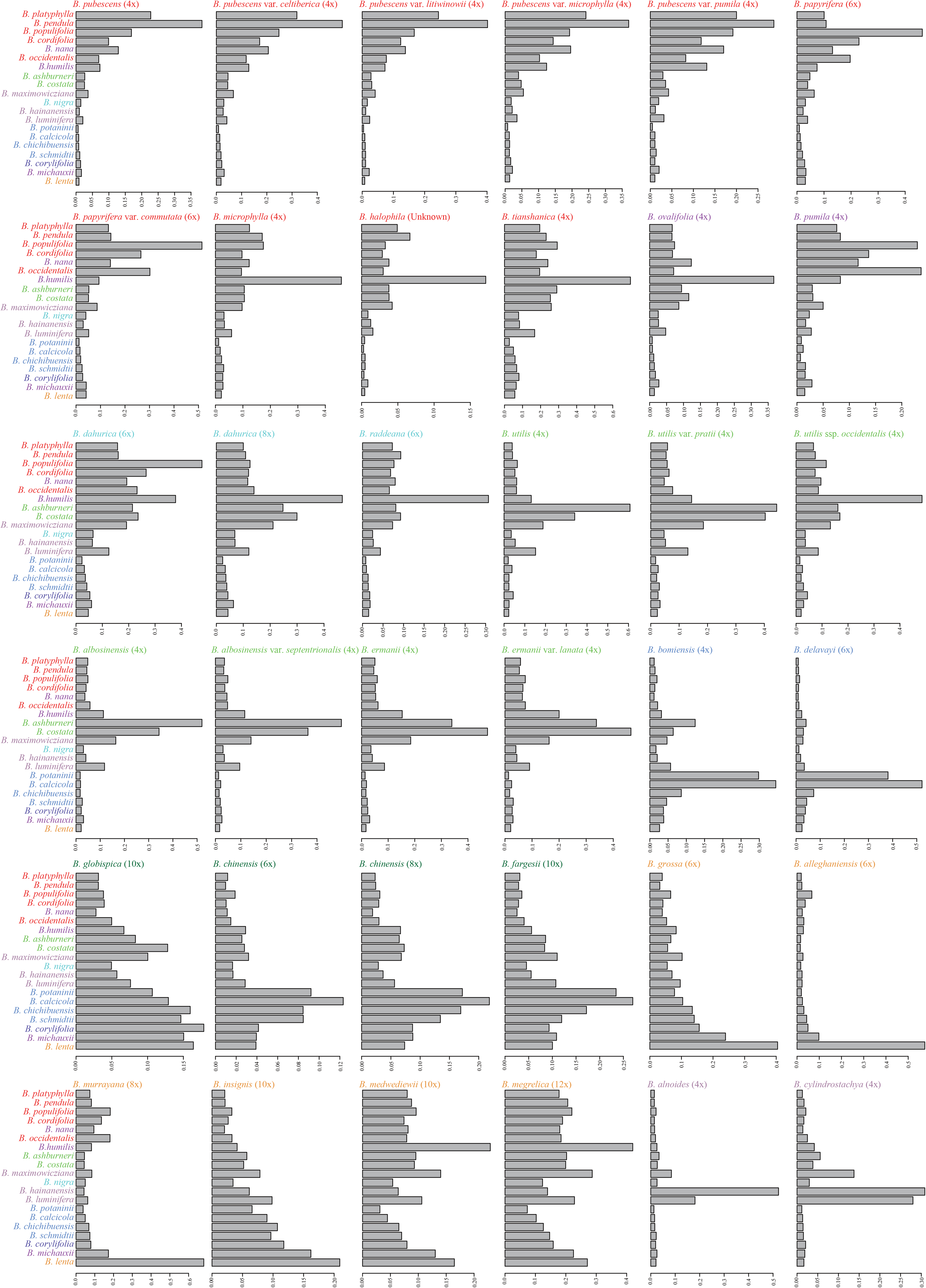
Mapping patterns of polyploids to the diploid reference. Numbers on the x axis indicate number of mapped loci.

### 3.5. Phylogeny incorporating polyploid species

When we included phased homoeologues from polyploids for which we could identify putative parents in a phylogenetic analysis, for 22 of the 26 the polyploids their homoeologues form clades with each of the putative parental diploid species (Table 1; Fig. 5). For example, subgenomes of *B. pubescens* and its varieties formed monophyletic clades which were sister to *B. pendula* and *B. platyphylla*, respectively.

**Figure 5.**
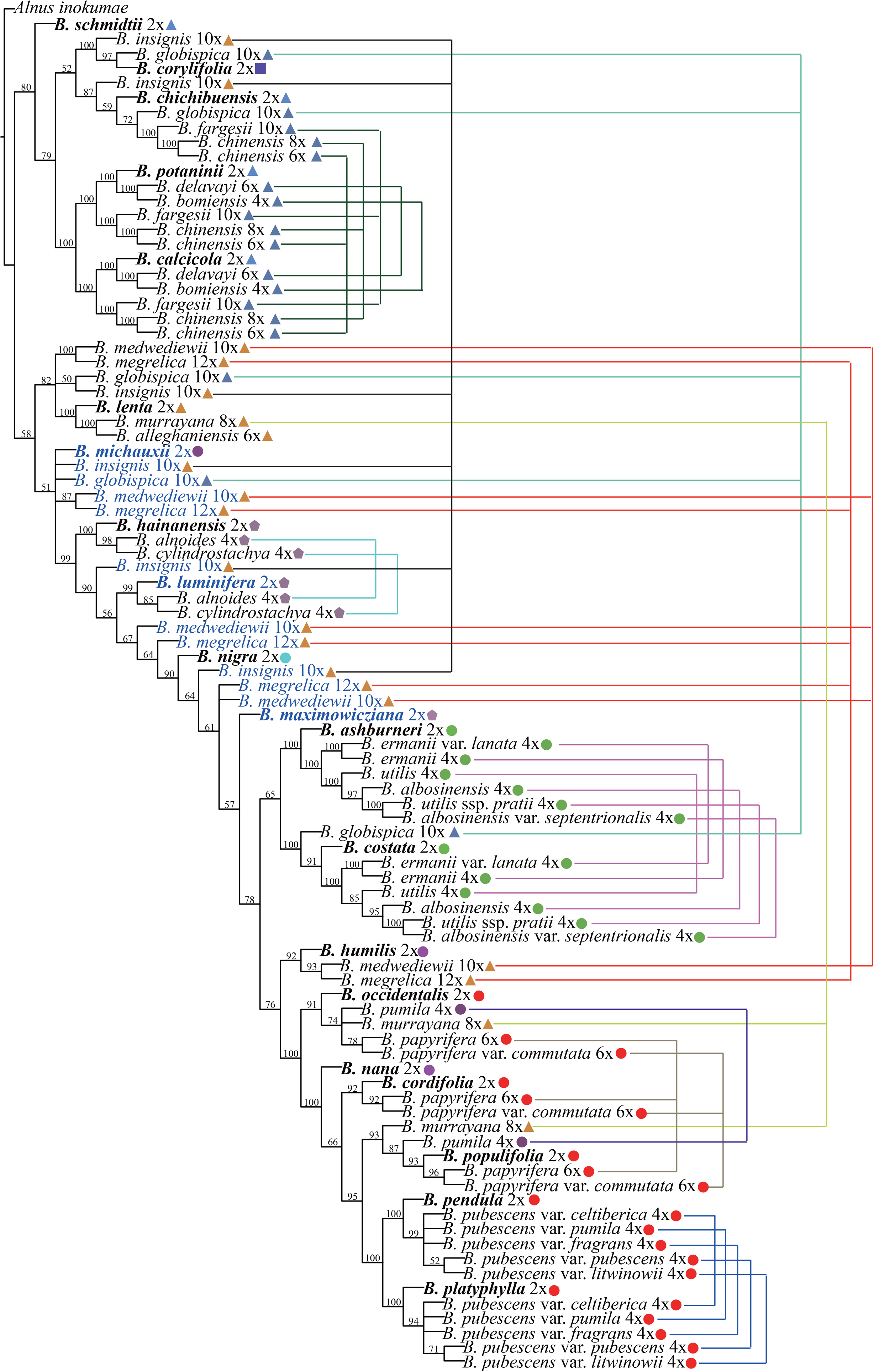
Species tree incorporating polyploids from the maximum likelihood analysis using ASTRID. Species were classified according to Ashburner and McAllister (2013).

### 3.6. Simple sequence repeat analysis

We developed between 58 and 565 microsatellite primer pairs for the diploid *Betula* taxa and between 40 and 633 for polyploid *Betula* taxa. In addition, 100, 84 and 41 microsatellite primers pairs were developed for *A. inokumae*, *A. orientalis* and *C. avellana*, respectively (Table S3).

## 4. Discussion

### 4.1. A well resolved diploid phylogeny for *Betula*

We used both supermatrix, and more unusually, species tree approaches to construct phylogenies based on RADseq data. We used longer contigs than is usual with RADseq (Tripp et al., 2017), with an average length of 675 bp; these contigs were generated from paired MiSeq reads, mapped to a reference genome (Salojarvi et al., 2017). The use of multiple gene trees also enabled us to detect evidence for hybridisation events among diploid species.

Two major clades were found in all analyses, which were not found by previous molecular or morphological analyses (Ashburner and McAllister, 2016; Bina et al., 2016; Järvinen et al., 2004; Li et al., 2005; Li et al., 2007; Nagamitsu et al., 2006; Schenk et al., 2008; Wang et al., 2016). Interestingly, species of Clade 1 exclusively have no or very narrow seed wings and species of Clade 2 exclusively have obvious seed wings. The fact that some species of Clade 2 have very wide geographic distributions is likely due to their strong dispersal ability. Ashburner and McAllister’s taxonomic sections *Asperae*, *Lentae* and *Nipponobetula* were grouped exclusively into Clade 1 and sections *Acuminatae*, *Dahuricae*, *Betula*, *Costatae*, were grouped exclusively into Clade 2, but species of section *Apterocaryon*, which are dwarf in form, are split between both clades. Within *Apterocaryon*, *B. michauxii*, with a geographic distribution in North America, is nested into Clade 1 whereas *B. humilis* and *B. nana*, with a geographic distribution across Eurasia, are nested within Clade 2. Thus the shrubby dwarf forms are likely due to independent evolution. Independent evolution of dwarf forms has been occasionally observed in other genera, such as in *Artemisia* (Tkach et al., 2007) and *Eucalyptus* (Foster et al., 2007). Another trait that may have evolved independently in the genus is resistance to the bronze birch borer: species reported to be largely resistant to the bronze birch borer (Muilenburg and Herms, 2012) are split between the two clades.

On the basis of the above, we suggest that section *Apterocaryon* should be dissolved, and *B. michauxii* placed in section *Lentae*, and *B. humilis* and *B. nana* in section *Betula*. In the case of section *Acuminatae*, our analysis shows *B. luminifera* and *B. haninanensis* form a clade, but *B. maximowicziana* is sister to section *Costatae*. The incongruence between morphology and molecular evidence for these three species is likely explained by hybridisation as indicated by our phylogenetic network analysis. We suggest that *B. maximowicziana* should be moved to section *Costatae*. These changes, taking into account the effects of hybridisation and convergent evolution, mean that the seven remaining sections of the genus *Betula*, and the three remaining subgenera proposed by Ashburner and McAllister, are all monophyletic in our diploid trees.

We also note that in our analyses, *B. corylifolia*, the single species of section *Nipponobetula*, was in a monophyletic group with species of section *Asperae*. Such a relationship has previously been indicated based on ITS (Nagamitsu et al., 2006; Wang et al., 2016). We therefore suggest that this species is placed in section *Asperae*, reducing the genus to two sub-genera that correspond to the two major clades of our diploid phylogenies. These are subgenus *Betula* containing sections *Acuminatae*, *Costatae, Dahuricae* and *Betula*, and subgenus *Aspera* containing sections *Asperae* and *Lentae*.

### 4.2. Inferring polyploid parentage of *Betula* species

Our results provide novel insights into parental species for a majority of *Betula* polyploids (Table 1). For example, tetraploids of section *Costatae* (excluding *B. utilis* ssp. *occidentalis*) have a high proportion of loci mapped to in *B. ashburneri* and *B. costata*, indicating their likely parentage. This is consistent with morphological characters, based on which Ashburner and McAllister placed these species within section *Costatae* (Ashburner and McAllister, 2016). The parentage of *B. pubescens*, which is widely planted, has been controversial for decades and has been suggested as *B. pendula* (Howland et al., 1995; Walters, 1968), *B. humilis*, *B. nana* (Howland et al., 1995; Järvinen et al., 2004; Walters, 1968) and *B. lenta* c.f. (Salojarvi et al., 2017). Here we find *B. pendula* and its sister species *B. platyphylla* to be the most likely parents of *B. pubescens*. We found a relatively low but still a considerable proportion of mapped loci from *B. pubescens* to *B. nana* and *B. humilis*, but several studies have found evidence for introgressive hybridisation among these species since the formation of *B. pubescens* (Bona et al., 2018; Jadwiszczak et al., 2012; Thórsson et al., 2001) which could account for sequences from *B. nana* and *B. humilis* in *B. pubsecens* genomes. Previous hypotheses for *B. lenta* c.f. as a parental species of *B. papyrifera* and *B. humilis* cf. as a parental species of *B. ermanii* (Järvinen et al., 2004) were not supported by our results. Our results also show evidence of complex layering of hybridisation and polyploidisation events in the history of some taxa. For example, *B. luminifera* and *B. hainanensis* were identified as the extant representatives for the progenitors of two tetraploids, but they themselves have a possible hybrid origin (or have been subject to significant introgression) on the basis of the network analysis; the two tetraploids with *B. luminifera*/*B. hainanensis* identified as progenitors both have the next highest proportion of loci mapped to in *B. maximowicziana*, which might suggest the evidence for hybridisation between *B. luminifera*/*B. hainanensis* and *B. maximowicziana* identified from the network analysis occurred before the origin of the polyploids. Given such complex evolutionary histories, it is perhaps unsurprising that for eight of the 35 polyploids, we could not clearly identify all putative parents. This may also be because these eight polyploids are older than the polyploids for which we have identified putative parents, and thus more divergent from the diploid species. Another possibility is that they are derived from diploid species which are now extinct, have not yet been discovered or were not included in our phylogenetic analyses (i.e. *B. glandulosa*).

## 5. Conclusion

Here, by generating a new phylogenetic hypothesis for *Betula*, and providing new evidence for the progenitors of many of its polyploid taxa, we have provided a framework within which the evolution and systematics of the genus can be understood. Knowledge of the parentage of the allopolyploids, some of which are widespread and economically important, opens the way for their genomic analysis. The approach we have used is relatively cheap and straightforward and could be applied to many other plant groups where allopolyploidy has impeded evolutionary analyses.

## Acknowledgements

This work was funded by Natural Environment Research Council Fellowship NE/G01504X/1 to R.J.A.B. and was funded by the National Natural Science Foundation of China (31770230, 31600295) to N.W.

## Author Contributions

NW and RB conceived the project. NW, RB and HM collected samples. HM identified the samples based on morphology. NW carried out lab work. NW, JZ and LK analysed data. NW, RB and LK wrote the manuscript. All the authors contributed to editing the manuscript.

---

**Figure S1** Mapping patterns of the 20 diploid *Betula* species of D1. Numbers on the x axis indicate number of mapped loci.

**Figure S2** The schematic illustration of methods used for analyzing polyploids.

**Figure S3** Species tree from the maximum likelihood analysis of the 20 *Betula* diploids using the ASTRID approach based on 50,870 gene trees. Species were classified according to Ashburner and McAllister (2016).

**Figure S4** Species tree from the maximum likelihood analysis of the 27 *Betula* diploids using the ASTRAL approach based on 58,442 gene trees. Asterisks on the branches indicate 1.00 local posterior probabilities. Species were classified according to Ashburner and McAllister (2016).

**Figure S5** Species tree from the maximum likelihood analysis of the 27 *Betula* diploids using the ASTRID approach based on 58,442 gene trees. Numbers above branches are support values. Marked with star indicates branches with low support values. Species were classified according to Ashburner and McAllister (2016).

**Figure S6** Species tree from the maximum likelihood analysis of the 27 *Betula* diploids using the supermatrix approach based on 58,442 gene trees. Numbers above or below branches are support values. The scale bar indicates the mean number of nucleotide substitutions per site. Species were classified according to Ashburner and McAllister (2016).

**Figure S7** The pseudolikelihood values for the number of hybridization events from 1 to 5.

## Supplementary data

**Table S1** Detailed information about *Betula* species used in this study and mapping results of RADseq.

**Table S2** Number of loci for each diploid species represented in the reference.

**Table S3** Detailed information about microsatellite markers mined from assembled contigs of species of *Betula*, *Alnus* and *Corylus*.

## Notes

### Competing Interest Statement

The authors have declared no competing interest.

## References

Anamthawat-Jónsson, K. and Tómasson, T., 1990. Cytogenetics of hybrid introgression in Icelandic birch. Hereditas 112, 65–70.

Anamthawat-Jónsson, K. and Tómasson, T., 1999. High frequency of triploid birch hybrid by *Betula nana* seed parent. Hereditas 130, 191–193.

Anamthawat-Jónsson, K. and Thórsson, A.T., 2003. Natural hybridisation in birch: triploid hybrids between *Betula nana* and *B. pubescens*. Plant Cell Tissue and Organ Culture 75, 99–107.

Anamthawat-Jónsson, K., Thórsson, A.T., Temsch, E.M. and Greilhuber, J., 2010. Icelandic birch polyploids-the case of perfect fit in genome size. Journal of Botany 347254.

Andrews, K.R., Good, J.M., Miller, M.R., Luikart, G. and Hohenlohe, P.A., 2016. Harnessing the power of RADseq for ecological and evolutionary genomics. Nature Reviews Genetics 17, 81–92.

Andrews, S., 2014. FastQC: a quality control tool for high throughput sequence data. http://www.bioinformatics.babraham.ac.uk/projects/fastqc/.

Ashburner, K. and McAllister, H.A., 2016. The Genus *Betula*: A Taxonomic Revision of Birches. Kew Publishing, London.

Avni, R., Nave, M., Barad, O., Baruch, K., Twardziok, S.O., Gundlach, H., Hale, I., Mascher, M., Spannagl, M., Wiebe, K., Jordan, K.W., Golan, G., Deek, J., Ben-Zvi, B., Ben-Zvi, G., Himmelbach, A., MacLachlan, R.P., Sharpe, A.G., Fritz, A., Ben-David, R., Budak, H., Fahima, T., Korol, A., Faris, J.D., Hernandez, A., Mikel, M.A., Levy, A.A., Steffenson, B., Maccaferri, M., Tuberosa, R., Cattivelli, L., Faccioli, P., Ceriotti, A., Kashkush, K., Pourkheirandish, M., Komatsuda, T., Eilam, T., Sela, H., Sharon, A., Ohad, N., Chamovitz, D.A., Mayer, K.F.X., Stein, N., Ronen, G., Peleg, Z., Pozniak, C.J., Akhunov, E.D. and Distelfeld, A., 2017. Wild emmer genome architecture and diversity elucidate wheat evolution and domestication. Science 357, 93–96.

Barchi, L., Lanteri, S., Portis, E., Acquadro, A., Valè, G., Toppino, L. and Rotino, G.L., 2011. Identification of SNP and SSR markers in eggplant using RAD tag sequencing. BMC Genomics 12, 304.

Barker, M.S., Arrigo, N., Baniaga, A.E., Li, Z. and Levin, D.A., 2016. On the relative abundance of autopolyploids and allopolyploids. New Phytol 210, 391–398.

Barnes, B.V., Bruce, P.D. and Sharik, T.L., 1974. Natural hybridization of yellow birch and white birch. Forest Science 20, 215–221.

Bina, H., Yousefzadeh, H., Ali, S.S. and Esmailpour, M., 2016. Phylogenetic relationships, molecular taxonomy, biogeography of *Betula*, with emphasis on phylogenetic position of Iranian populations. Tree Genet Genomes 12, 84.

Bolger, A.M., Lohse, M. and Usadel, B., 2014. Trimmomatic: a flexible trimmer for Illumina sequence data. Bioinformatics 30, 2114–2120.

Bona, A., Petrova, G. and Jadwiszczak, K.A., 2018. Unfavourable habitat conditions can facilitate hybridisation between the endangered *Betula humilis* and its widespread relatives *B. pendula* and *B. pubescens*. Plant Ecology & Diversity, doi:10.1080/17550874.17552018.11518497.

Brysting, A.K., Mathiesen, C. and Marcussen, T., 2011. Challenges in polyploid phylogenetic reconstruction: a case story from the arctic-alpine Cerastium alpinum complex. Taxon 60, 333–347.

Capella-Gutierrez, S., Silla-Martinez, J.M. and Gabaldon, T., 2009. trimAl: a tool for automated alignment trimming in large-scale phylogenetic analyses. Bioinformatics 25, 1972–1973.

Cariou, M., Duret, L. and Charlat, S., 2013. Is RAD-seq suitable for phylogenetic inference? An in silico assessment and optimization. Ecology and Evolution 3, 846–852.

Catchen, J., Hohenlohe, P.A., Bassham, S., Amores, A. and Cresko, W.A., 2013. Stacks: an analysis tool set for population genomics. Mol Ecol 22, 3124–3140.

Cruaud, A., Gautier, M., Galan, M., Foucaud, J., Saune, L., Genson, G., Dubois, E., Nidelet, S., Deuve, T. and Rasplus, J.Y., 2014. Empirical assessment of RAD sequencing for interspecific phylogeny. Molecular Biology and Evolution 31, 1272–1274.

DaCosta, J.M. and Sorenson, M.D., 2016. ddRAD-seq phylogenetics based on ucleotide, indel, and presence-absence polymorphisms: Analyses of two avian genera with contrasting histories. Mol Phylogenet Evol 94, 122–135.

Eaton, D.A.R. and Ree, R.H., 2013. Inferring Phylogeny and Introgression using RADseq Data: An Example from Flowering Plants (*Pedicularis*: Orobanchaceae). Syst Biol 62, 689–706.

Eaton, D.A.R., Spriggs, E.L., Park, B. and Donoghue, M.J., 2016. Misconceptions on missing data in RAD-seq phylogenetics with a deep-scale example from flowering plants. Syst Biol 16, syw092.

Eidesen, P.B., Alsos, I.G. and Brochmann, C., 2015. Comparative analyses of plastid and AFLP data suggest different colonization history and asymmetric hybridization between *Betula pubescens* and *B. nana*. Mol Ecol 24, 3993–4009

Emerson, K.J., Merz, C.R., Catchen, J.M., Hohenlohe, P.A., Cresko, W.A., Bradshaw, W.E. and Holzapfel, C.M., 2010. Resolving postglacial phylogeography using high-throughput sequencing. Proceedings of the National Academy of Sciences of the United States of America 107, 16196–16200.

Eriksson, J.S., de Sousa, F., Bertrand, Y.J.K., Antonelli, A., Oxelman, B. and Pfeil, B.E., 2018. Allele phasing is critical to revealing a shared allopolyploid origin of *Medicago arborea* and *M. strasseri* (Fabaceae). Bmc Evol Biol 18, 9.

Etter, P.D., Bassham, S., Hohenlohe, P.A., Johnson, E.A. and Cresko, W.A., 2011. SNP discovery and genotyping for evolutionary genetics using RAD sequencing. In: Orgogonzo, V., Rockman, M.V. (Eds.), Molecular Methods for Evolutionary Genetics. Humana Press, NY.

Folk, R.A., Mandel, J.R. and Freudenstein, J.V., 2017. Ancestral Gene Flow and Parallel Organellar Genome Capture Result in Extreme Phylogenomic Discord in a Lineage of Angiosperms. Syst Biol 66, 320–337.

Fontaine, M.C., Pease, J.B., Steele, A., Waterhouse, R.M., Neafsey, D.E., Sharakhov, I.V., Jiang, X.F., Hall, A.B., Catteruccia, F., Kakani, E., Mitchell, S.N., Wu, Y.C., Smith, H.A., Love, R.R., Lawniczak, M.K., Slotman, M.A., Emrich, S.J., Hahn, M.W. and Besansky, N.J., 2015. Extensive introgression in a malaria vector species complex revealed by phylogenomics. Science 347.

Foster, S.A., McKinnon, G.E., Steane, D.A., Potts, B.M. and Vaillancourt, R.E., 2007. Parallel evolution of dwarf ecotypes in the forest tree *Eucalyptus globulus*. New Phytol 175, 370–380.

Gonen, S., Bishop, S.C. and Houston, R.D., 2015. Exploring the utility of cross-laboratory RAD-sequencing datasets for phylogenetic analysis. BMC Research Notes 8, 299.

Hipp, A.L., Eaton, D.A.R., Cavender-Bares, J., Fitzek, E., Nipper, R. and Manos, P.S., 2014. A Framework Phylogeny of the American Oak Clade Based on Sequenced RAD Data. Plos One 9.

Hohenlohe, P.A., Bassham, S., Etter, P.D., Stiffler, N., Johnson, E.A. and Cresko, W.A., 2010. Population genomics of parallel adaptation in threespine stickleback using sequenced RAD tags. PLoS Genetics 6, e1000862.

Hou, Y., Nowak, M.D., Mirre, V., Bjora, C.S., Brochmann, C. and Popp, M., 2015. Thousands of RAD-seq Loci Fully Resolve the Phylogeny of the Highly Disjunct Arctic-Alpine Genus *Diapensia* (Diapensiaceae). Plos One 10, e0140175.

Howland, D.E., Oliver, R.R. and Davy, A.J., 1995. Morphological and molecular variation in natural populations of *Betula*. New Phytol 130, 117–124.

Järvinen, P., Palmé, A., Morales, L.O., Lännenpää, M., Keinänen, M., Sopanen, T. and Lascoux, M., 2004. Phylogenetic relationships of *Betula* species (Betulaceae) based on nuclear ADH and chloroplast matK sequences. American Journal of Botany 91, 1834–1845.

Jadwiszczak, K.A., Banaszek, A., Jabłońska, E. and Sozinov, O.V., 2012. Chloroplast DNA variation of *Betula humilis* Schrk. in Poland and Belarus. Tree Genet Genomes 8, 1017–1030.

Johnsson, H., 1945. Interspecific hybridization within the genus *Betula*. Hereditas 31, 163–176.

Jones, G., Sagitov, S. and Oxelman, B., 2013. Statistical Inference of Allopolyploid Species Networks in the Presence of Incomplete Lineage Sorting. Syst Biol 62, 467–478.

Katoh, K., Kuma, K., Toh, H. and Miyata, T., 2005. MAFFT version 5: improvement in accuracy of multiple sequence alignment. Nucleic Acids Research 33, 511–518.

Li, F.G., Fan, G.Y., Lu, C.R., Xiao, G.H., Zou, C.S., Kohel, R.J., Ma, Z.Y., Shang, H.H., Ma, X.F., Wu, J.Y., Liang, X.M., Huang, G., Percy, R.G., Liu, K., Yang, W.H., Chen, W.B., Du, X.M., Shi, C.C., Yuan, Y.L., Ye, W.W., Liu, X., Zhang, X.Y., Liu, W.Q., Wei, H.L., Wei, S.J., Huang, G.D., Zhang, X.L., Zhu, S.J., Zhang, H., Sun, F.M., Wang, X.F., Liang, J., Wang, J.H., He, Q., Huang, L.H., Wang, J., Cui, J.J., Song, G.L., Wang, K.B., Xu, X., Yu, J.Z., Zhu, Y.X. and Yu, S.X., 2015. Genome sequence of cultivated Upland cotton (*Gossypium hirsutum* TM-1) provides insights into genome evolution. Nat Biotechnol 33, 524–530.

Li, G., Davis, B.W., Eizirik, E. and Murphy, W.J., 2016. Phylogenomic evidence for ancient hybridization in the genomes of living cats (Felidae). Genome Res 26, 1–11.

Li, J.H., Shoup, S. and Chen, Z.D., 2005. Phylogenetics of *Betula* (Betulaceae) inferred from sequences of nuclear ribosomal DNA. Rhodora 107, 69–86.

Li, J.H., Shoup, S. and Chen, Z.D., 2007. Phylogenetic relationships of diploid species of *Betula* (Betulaceae) inferred from DNA sequences of nuclear nitrate reductase. Systematic Botany 32, 357–365.

Linder, C.R. and Rieseberg, L.H., 2004. Reconstructing patterns of reticulate evolution in plants. American Journal of Botany 91, 1700–1708.

Lott, M., Spillner, A., Huber, K.T., Petri, A., Oxelman, B. and Moulton, V., 2009. Inferring polyploid phylogenies from multiply-labeled gene trees. Bmc Evol Biol 9, 216.

Luo, M.C., Gu, Y.Q., Puiu, D., Wang, H., Twardziok, S.O., Deal, K.R., Huo, N.X., Zhu, T.T., Wang, L., Wang, Y., McGuire, P.E., Liu, S.Y., Long, H., Ramasamy, R.K., Rodriguez, J.C., Van, S.L., Yuan, L.X., Wang, Z.Z., Xia, Z.Q., Xiao, L.C., Anderson, O.D., Ouyang, S.H., Liang, Y., Zimin, A.V., Pertea, G., Qi, P., Ennetzen, J.L.B., Dai, X.T., Dawson, M.W., Muller, H.G., Kugler, K., Rivarola-Duarte, L., Spannagl, M., Mayer, K.F.X., Lu, F.H., Bevan, M.W., Leroy, P., Li, P.C., You, F.M., Sun, Q.X., Liu, Z.Y., Lyons, E., Wicker, T., Salzberg, S.L., Devos, K.M. and Dvorak, J., 2017. Genome sequence of the progenitor of the wheat D genome *Aegilops tauschii*. Nature 551, 498–502.

Massatti, R., Reznicek, A.A. and Knowles, L.L., 2016. Utilizing RADseq data for phylogenetic analysis of challenging taxonomic groups: A case study in *Carex* sect. *Racemosae*. American Journal of Botany 103, 337–347.

McKain, M.R., Tang, H., McNeal, J.R., Ayyampalayam, S., Davis, J.I., depamphilis, C.W., Givnish, T.J., Pires, J.C., Stevenson, D.W. and Leebens-Mack, J.H., 2016. A Phylogenomic Assessment of Ancient Polyploidy and Genome Evolution across the Poales. Genome Biol Evol 8, 1150–1164.

Meglécz, E., Pech, N., Gilles, A., Dubut, V., Hingamp, P., Trilles, A. and Grenier, R.e.a., 2014. QDD version 3.1: a user-friendly computer program for microsatellite selection and primer design revisited: experimental validation of variables determining genotyping success rate. Mol Ecol Resour 14, 1302–1313.

Mirarab, S. and Warnow, T., 2015. ASTRAL-II: coalescent-based species tree estimation with many hundreds of taxa and thousands of genes. Bioinformatics 31, i44–i52.

Morales-Briones, D.F., Liston, A. and Tank, D.C., 2018. Phylogenomic analyses reveal a deep history of hybridization and polyploidy in the Neotropical genus *Lachemilla* (Rosaceae). New Phytol 218, 1668–1684.

Muilenburg, V.L. and Herms, D.A., 2012. A Review of Bronze Birch Borer (Coleoptera: Buprestidae) Life History, Ecology, and Management. Environ Entomol 41, 1372–1385.

Nagamitsu, T., Kawahara, T. and Kanazashi, A., 2006. Endemic dwarf birch *Betula apoiensis* (Betulaceae) is a hybrid that originated from *Betula ermanii* and *Betula ovalifolia*. Plant Species Biology 21, 19–29.

Oxelman, B., Brysting, A.K., Jones, G.R., Marcussen, T., Oberprieler, C. and Pfeil, B.E., 2017. Phylogenetics of Allopolyploids. Annual Review of Ecology, Evolution, and Systematics 48, 543–557.

Pante, E., Abdelkrim, J., Viricel, A., Gey, D., France, S.C., Boisselier, M.C. and Samadi, S., 2015. Use of RAD sequencing for delimiting species. Heredity 114, 450–459.

Rothfels, C.J., Pryer, K.M. and Li, F.W., 2017. Next-generation polyploid phylogenetics: rapid resolution of hybrid polyploid complexes using PacBio single-molecule sequencing. New Phytol 213, 413–429.

Rubin, B.E.R., Ree, R.H. and Moreau, C.S., 2012. Inferring phylogenies from RAD sequence data. Plos One 7, e33394.

Salojarvi, J., Smolander, O.P., Nieminen, K., Rajaraman, S., Safronov, O., Safdari, P., Lamminmaki, A., Immanen, J., Lan, T.Y., Tanskanen, J., Rastas, P., Amiryousefi, A., Jayaprakash, B., Kammonen, J.I., Hagqvist, R., Eswaran, G., Ahonen, V.H., Serra, J.A., Asiegbu, F.O., Barajas-Lopez, J.D., Blande, D., Blokhina, O., Blomster, T., Broholm, S., Brosche, M., Cui, F.Q., Dardick, C., Ehonen, S.E., Elomaa, P., Escamez, S., Fagerstedt, K.V., Fujii, H., Gauthier, A., Gollan, P.J., Halimaa, P., Heino, P.I., Himanen, K., Hollender, C., Kangasjarvi, S., Kauppinen, L., Kelleher, C.T., Kontunen-Soppela, S., Koskinen, J.P., Kovalchuk, A., Karenlampi, S.O., Karkonen, A.K., Lim, K.J., Leppala, J., Macpherson, L., Mikola, J., Mouhu, K., Mahonen, A.P., Niinemets, U., Oksanen, E., Overmyer, K., Palva, E.T., Pazouki, L., Pennanen, V., Puhakainen, T., Poczai, P., Possen, B.J.H.M., Punkkinen, M., Rahikainen, M.M., Rousi, M., Ruonala, R., van der Schoot, C., Shapiguzov, A., Sierla, M., Sipila, T.P., Sutela, S., Teeri, T.H., Tervahauta, A.I., Vaattovaara, A., Vahala, J., Vetchinnikova, L., Welling, A., Wrzaczek, M., Xu, E.J., Paulin, L.G., Schulman, A.H., Lascoux, M., Albert, V.A., Auvinen, P., Helariutta, Y. and Kangasjarvi, J., 2017. Genome sequencing and population genomic analyses provide insights into the adaptive landscape of silver birch. Nat Genet 49, 904–912.

Sayyari, E. and Mirarab, S., 2016. Fast coalescent-based computation of local branch support from quartet frequencies. Molecular Biolology and Evolution 33, 1654–1668.

Schenk, M.F., Thienpont, C.N., Koopman, W.J.M., Gilissen, L.J.W.J. and Smulders, M.J.M., 2008. Phylogenetic relationships in Betula (Betulaceae) based on AFLP markers. Tree Genet Genomes 4, 911–924.

Shaw, K., Stritch, L., Rivers, M., Roy, S., Wilson, B. and Govaerts, R., 2014. The Red List of Betulaceae. BGCI, Richmond. UK.

Solís-Lemus, C. and Ané, C., 2016. Inferring phylogenetic networks with maximum pseudolikelihood under incomplete lineage sorting. PLoS Genetics 12, e1005896.

Solis-Lemus, C., Bastide, P. and Ane, C., 2017. PhyloNetworks: A Package for Phylogenetic Networks. Mol Biol Evol 34, 3292–3298.

Stamatakis, A., 2006. RAxML-VI-HPC: maximum likelihood-based phylogenetic analyses with thousands of taxa and mixed models. Bioinformatics 22, 2688–2690.

Thórsson, T.H., Salmela, E. and Anamthawat-Jónsson, K., 2001. Morphological, cytogenetic, and molecular evidence for introgressive hybridization in birch. Journal of Heredity 92, 404–408.

Tkach, N.V., Hoffmann, M.H., Röser, M., Korobkov, A.A. and von Hagen, K.B., 2007. Parallel evolutionary patterns in multiple lineages of arctic *Artemisia* L. (Asteraceae). Evolution 62, 184–198.

Tripp, E.A., Tsai, Y.H.E., Zhuang, Y. and Dexter, K.G., 2017. RADseq dataset with 90% missing data fully resolves recent radiation of *Petalidium* (Acanthaceae) in the ultra-arid deserts of *Namibia*. Ecology and Evolution 7, 7920–7936.

Tsuda, Y., Semerikov, V., Sebastiani, F., Vendramin, G.G. and M., L., 2017. Multispecies genetic structure and hybridization in the *Betula* genus across Eurasia. Mol Ecol 26, 589–605

Untergasser, A., Cutcutache, I., Koressaar, T., Ye, J., Faircloth, B.C., Remm, M. and Rozen, S.G., 2012. Primer3-new capabilities and interfaces. Nucleic Acids Res 40, e115.

Vachaspati, P. and Warnow, T., 2015. ASTRID: Accurate Species TRees from Internode Distances. Bmc Genomics 16.

Wagner, C.E., Keller, I., Wittwer, S., Selz, O.M., Mwaiko, S., Greuter, L., Sivasundar, A. and Seehausen, O., 2013. Genome-wide RAD sequence data provide unprecedented resolution of species boundaries and relationships in the Lake Victoria cichlid adaptive radiation. Mol Ecol 22, 787–798.

Walters, S.M., 1968. *Betula* L. in Britain. Proceedings of the Botanical Society of the British Isles 7, 179–180.

Wang, N., Borrell, J.S., Bodles, W.J.A., Kuttapitiya, A., Nichols, R.A. and Buggs, R.J.A., 2014. Molecular footprints of the Holocene retreat of dwarf birch in Britain. Mol Ecol 23, 2771–2782.

Wang, N., McAllister, H.A., Bartlett, P.R. and Buggs, R.J.A., 2016. Molecular phylogeny and genome size evolution of the genus *Betula* (Betulaceae). Ann Bot-London 117, 1023–1035.

Wang, N., Thomson, M., Bodles, W.J.A., Crawford, R.M.M., Hunt, H.V., Featherstone, A.W., Pellicer, J. and Buggs, R.J.A., 2013. Genome sequence of dwarf birch (*Betula nana*) and cross-species RAD markers. Mol Ecol 22, 3098–3111.

Zohren, J., Wang, N., Kardailsky, I., Borrell, J.S., Joecker, A., Nichols, R.A. and Buggs, R.J.A., 2016. Unidirectional diploid-tetraploid introgression among British birch trees with shifting ranges shown by restriction site-associated markers. Mol Ecol 25, 2413–2426.

